# Involvement of multiple influx and efflux transporters in the accumulation of cationic fluorescent dyes by *Escherichia coli*

**DOI:** 10.1101/603688

**Authors:** Srijan Jindal, Lei Yang, Philip J. Day, Douglas B. Kell

## Abstract

We used high-throughput flow cytometry to assess the ability of individual gene knockout strains of *E coli* to take up two membrane-permeable, cationic fluorescent dyes, viz the carbocyanine diS-C3(5) and the DNA dye SYBR Green. Individual strains showed a large range of distributions of uptake. The range of modal steady-state uptakes for the carbocyanine between the different strains was 36-fold. Knockouts of the ATP synthase α- and β-subunits greatly inhibited uptake, implying that most uptake was ATP-driven rather than being driven by say a membrane potential. Dozens of transporters changed the steady-state uptake of the dye by more than 50% with respect to that of the wild type, in both directions (increased or decreased); knockouts in known influx and efflux transporters behaved as expected, giving confidence in the general strategy. Many of the knockouts with the most reduced uptake were transporter genes of unknown function (‘y-genes’). Similarly, several overexpression variants in the ‘ASKA’ collection had the anticipated, opposite effects. Similar findings were made with SYBR Green (the range being some 69-fold), though despite it too containing a benzimidazole motif there was negligible correlation between its uptake and that of the carbocyanine when compared across the various strains. Overall, we conclude that the uptake of these dyes may be catalysed by a great many transporters of possibly broad and presently unknown specificity. This casts serious doubt upon the use of such dyes as quantitative stains for representing either bioenergetic parameters or the amount of cellular DNA in unfixed cells (*in vivo*). By contrast, it opens up their potential use as transporter assay substrates in high-throughput screening.

## Introduction

The presence, number, nature, and physiological status of bacteria is widely assessed using fluorescent dyes [1-9]. Some of these stain particular molecules or macromolecules such as DNA [10], protein [11] or lipid [12], while others reflect physiological variables such as pH, the concentrations of other ions and small molecules, or the extent of membrane energisation [13]. In some cases (e.g. [6; 8]), cells are permeabilised before staining, such that the natural ability or otherwise of stains to reach their intracellular targets is not an issue. However, studies of *in vivo* physiology [9; 14-16] necessarily require that native, intact cells are used. In Gram-negative cells, entry to the periplasm is mediated via porins [17], which can affect the ability of stains such as rhodamine 123 and other cations to be accumulated [18-20], while intracellular concentrations of stain may also be modified by the activities of influx and efflux transporters.

Although it is often assumed and stated that intact cells take up xenobiotics mainly according to their hydrophobicity (lipophilicity) [21-23], and generally by passage through the limited phospholipid bilayer areas that exist, an abundance of evidence indicates that this is not in fact the case [24-26] and that ‘phospholipid bilayer transport is negligible’ [27]. Those xenobiotics that pass through the membrane, <u>which necessarily include xenobiotic fluorescent probes</u>, must thus be taken up by protein-based transporters, and the question arises as to which ones [28; 29]. In mammalian cells, for instance, fluorescein can be transported by an active monocarboxylate transporter [30].

Since the amount of fluorescence is often used as a quantitative indicator of the amount of the relevant determinand, it is obvious that the involvement of such transporters might significantly obscure the true values that would result if transporter-mediated membrane permeability or translocation was not occurring or was kinetically irrelevant. An early indication of this was the recognition that the apparent failure of ethidium bromide to stain intracellular DNA inside intact (live) *E. coli* cells was almost entirely due to the overwhelming activity of an efflux pump whose activity was much reduced at 0°C [31]. The relevant efflux pumps were not then identified, but it is now well established that ethidium bromide is a very good substrate for ‘efflux’ transporters such as the multisubunit acrAB-tolC complex [32] and the small molecule resistance (SMR) protein emrE [33].

Concentrative transporters (whether influx or efflux) need additional sources of free energy. In *E. coli* these are typically the energised membrane (often seen as a membrane potential or a protonmotive force [34]) generated via electron transport (that can also be used to synthesise ATP), or ATP itself acting directly. The latter is significantly more common in prokaryotes [35]. However, it is often considered that the uptake of more or less lipophilic cations into bacteria or mitochondria is driven by a negative-inside membrane potential, whose concentration ratio reflects it according to the Nernst equation and thereby allows one to infer it (see Discussion).

Many non-fluorescent drugs and detergents, including anti-infectives, can simply be seen as xenobiotics, and systematic studies have been performed to see the extent to which the loss of effluxers (and occasionally of influxers) modulates their toxicity [36; 37]. In particular, the AcrAB-TolC complex spans inner and outer membranes, is constitutively expressed, and is considered to play a major role in multidrug resistance [38-42]. Consequently, the activities of efflux transporters are widely recognised both as major mediators of microbial resistance to antibiotics and as targets for ameliorating it [43-69]. However, such activities cannot yet be predicted reliably (e.g. [70-72]). Efflux transporters are also important in pharmacokinetics, not least by effecting the export of anticancer drugs in mammalian systems (e.g. [53; 73-76]). They can also play important roles in the biotechnological production of small molecules and/or their biotransformations [77; 78].

Since efflux transporters are required to remove such intracellular molecules that have been taken up by cells, it is reasonable that influx transporters were exploited to get them there in the first place [24-27]. However, in the case of bacteria, knowledge of the specific influx and efflux transporters for individual xenobiotics is surprisingly limited, albeit many efflux transporters can be quite promiscuous [79-81].

Although they can be interfered with by coloured compounds [82], an understanding of the transporters used in the uptake and efflux of fluorescent probes in intact microbial cells is of interest for a number of reasons: (i) as with mammalian cells [83-86], they can provide substrates for competitive or trans-stimulation-type uptake assays, (ii) they provide examples of substrates that can be used in the development of quantitative structure-activity relationships (QSARs) for the effluxers themselves, and (iii) they allow us to assess the limitations of any individual fluorescent probe assay where the expression levels of relevant transporters is not known or controlled [87]. It is already known that even lipid stains such as Nile Red [88-91] and membrane energisation stains such as bis-(1,3-dibutylbarbituric acid trimethine oxonol) (commonly known as DiBAC4(3)) are in fact efflux substrates of acrAB-tolC [92], while other widely used stains that are effluxed via acrAB-tolC include the Hoechst dye H33342 [93], berberine [94], resazurin [95], and rhodamine 6G [96]. What is much less well known is which if any other efflux transporters are involved, and which influx transporters may have been responsible for the uptake of such commonly used fluorophores in the first place.

Following the systematic genome sequencing of *E. coli* K12 strain MG1655 [97], and an equivalent programme in baker’s yeast [98; 99], it was soon recognised that much scientific value would accrue to the possession of a collection of single-gene knockouts (of ‘non-essential’ genes), and this was produced as the ‘Keio collection’ [100-102] http://ecoli.iab.keio.ac.jp/. Flow cytometry provides a convenient means of estimating the steady-state uptake of fluorescent probes in bacteria [9], and while efflux pumps have been analysed elegantly in this way [44] the combination of flow cytometry and the Keio collection seemed to provide an ideal opportunity to assess the contribution of individual transporter genes (i.e. their products) to the uptake <u>and</u> efflux of widely used fluorescent probes. We used TransportdB http://www.membranetransport.org/transporter2.php?oOID=ecol1 [103] to pick out the most pertinent subset of transporters, to which we added a few more strains whose knockouts were involved in ATP synthesis, and report the findings herein. It is concluded that every probe used can exploit a wide variety of transporters for both influx and efflux, additional to reporting on their nominal determinand of interest. This is consistent with the view [27; 29; 104] that most xenobiotics that enter cells can be taken up and/or effluxed by multiple transporters of varying specificities.

## Materials and methods

### Bacterial strains

*E. coli* (K-12, MG1655) was taken from the laboratory collection of Prof R. Goodacre [105; 106]). The Keio collection of *E. coli* (K-12, MG1655) was obtained from the SYNBIOChem group (University of Manchester, UK) from which a collection of 530 knockouts (188 y-genes) was selected for the study. Only the strains that had a transporter protein gene knocked out were selected for the study. A few strains were selected from the ASKA collection of *E. coli* (BW38029) also taken from SYNBIOChem group.

### Culture

*E. coli* strains were grown from single colonies on agar plates in conical flasks using Lysogeny broth (LB) to an optical density (600 nm) of 1.5–2.0, representing stationary phase in this medium. They were held in stationary phase for 2–4 h before being inoculated at a concentration of 10^5^ cells.mL^-1^ into LB.

### Keio collection sample preparation

Singer ROTOR HDA (Singer Instruments, UK), a high throughput robotic replicator and colony picker, was used to pick single strains from stock 96-well plates stored at −80°C and stamped on nutrient agar plates in 96-well format. The agar plates were grown overnight and strains were then transferred in a 96-well plate with 200uL of LB in each well. The resulting set of six 96-well plates was incubated overnight at 37°C and 200 rpm shaking and used later for analysis.

### DiSC3(5) and Sybr Green I uptake measurement

Singer RePads 96 long (Singer Instruments, UK), 96-pin plastic replica-plating pads were used to replicate and culture the new subset of the Keio collection grown overnight. 150uL of LB was added in each well with 3uM (final concentration) of DiSC3(5) (Thermo Fisher Scientific, UK) in six U-bottom 96-well plates. The Singer RePads were used to transfer individual strains from overnight cultured plates to the new plates. The plates were sealed with Breathable Film (Starlab UK, Ltd.) and incubated at 37°C for 2 minutes in the dark. The plates were vortexed at 200 rpm for 2 min and screened on a Sartorius Intellicyt iQue Screener Plus™ flow cytometer.

Our variant of this flow cytometer has three fixed excitation lasers (405, 488, 640nm), forward and side scatter (from the 488nm excitation) and 13 fluorescence channels based on filters. The channels we report here are mainly BL1 (Ex 488, Em 530 ± 15nm) and RL1 (Ex 640nm, Em 675 ± 15nm). To detect bacteria we gated via forward scatter and side scatter. This instrument does not have user-adjustable photomultiplier tubes so the numbers simply reflect the extent of fluorescence it registers. The following settings [107] were used for the flow cytometry: automatic prime – 60 s (in Qsol buffer); pre-plate shake –15 s and 900 r.p.m.; sip time –2 s (actual sample uptake); additional sip time –0.5 s (the gap between sips); pump speed –29 r.p.m. (1.5 µl.s^−1^ sample uptake); plate model – U-bottom well plate (for 96-well plates); mid-plate clean-up – after every well (4 washes; 0.5 s each in Qsol buffer); inter-well shake –900 rpm; after 11 wells, 4 s in Qsol buffer; flush and clean – 30 s with Decon and Clean buffers followed by 60 s with de-ionized water. Fluorescence intensity was measured with RL1 channel and the Forecyt™ software supplied with the instrument was used to perform the analysis.

For Sybr Green I (Thermo Fisher Scientific, UK) uptake measurements, the samples were prepared similarly to those for DiSC3(5) except that Sybr Green I was added at 10,000x diluted concentration from the original stock. The sample plates were incubated for 15 minutes and fluorescence was measured using the BL1 channel.

The fluorescence kinetics experiment was performed on a Sony SH800 FACS machine. An overnight-grown culture of *E. coli* was diluted in LB to a concentration of 105 CFU.mL^-1^. diSC3(5) dye was added to 10mL of the bacterial solution at a final concentration of 3μM in a 50mL amber falcon tube (to prevent photobleaching) and incubated in the dark at 37°C for 2 minutes. Post-vortexing, the falcon tube was placed in the flow cytometer and the fluorescence intensity of the sample was measured for 1 hour continuously. Same was repeated with SYBR Green I.

### SYBR Green I uptake measurement on fixed Keio strains

A further subset of 20 strains from the previously analysed Keio collection subset was taken and fixed by injection in 70% ice-cold ethanol. The cells were washed twice by centrifugation in 0.1 M-Tris/HCl buffer, pH 7.4, before being resuspended in PBS at the concentration of 10^6^ CFU.mL^-1^. Sybr Green I was added to the sample at 10,000× diluted concentration from the stock and the samples were kept at 37°C in the dark for 15 minutes. Samples were added in a U-bottom 96-well plate and fluorescence intensity was measured using the BL1 channel.

### Effect of inhibitors on the fluorescence of dye uptake in *E. coli*

A few efflux inhibitors were tested for their effects on the fluorescence intensity of the bacterial cells. The stock solutions of the inhibitors were made at 1mM in DMSO and 5uL was added in two 96-well plates in triplicates. A vacuum centrifuge was used to evaporate all the DMSO in the plates. 200uL of overnight-cultured *E. coli* was added in each well of the plates at 10^5^ CFU.mL^-1^ final concentrations. The plates were sealed and kept at 30°C and 900rpm shaking for 30 minutes. 3uM of DiSC3(5) and 10,000× diluted Sybr Green I was then added to the different plates. The plate with DiSC3(5) was incubated for 2 minutes while one with Sybr Green I was incubated for 15 minutes at 37°C before being analysed using the flow cytometer.

### Screening ASKA collection strains for dye uptake

Selected strains from the ASKA collection were taken from −80°C stock and resuscitated in LB with 34 μg.mL^-1^ Chloramphenicol. The strains were cultured overnight in M9 media (M9 Minimal Media: 1× M9 Salts, 2 mM MgSO_4_, 0.1 mM CaCl_2_, 0.2% glucose, 10 ug.mL^-1^ Thiamine HCl and 0.2% Casamino Acids) and 0.5 mM Isopropyl β-D-1-thiogalactopyranoside (IPTG) was added in the morning and grown for another hour. The strains were diluted and their fluorescence upon uptake of DiSC3(5) was measured using the same steps as performed above with the Keio strains.

### Growth assessment of Keio collection strains by bulk OD measurement

The Growth Profiler 960 (Enzyscreen, NL) is a commercial instrument (http://www.enzyscreen.com/growth_profiler.htm) that estimates growth rates using camera-based measurements in up to 10 96-well plates. *E. coli* was taken from an overnight culture and diluted to a concentration of 10^6^ CFU.mL^-1^. The Keio collection strains were sub-cultured and samples were prepared using stamping method as described above. The strains were stamped in CR1496c Polystyrene transparent square 96-half-deepwell microtiter plates used with CR1396b Sandwich covers in the Growth Profiler. The samples were prepared in 10 96-well plates and incubated in the instrument at 37°C with 225 rpm shaking (recommended settings). The pictures of the plates were recorded at 15-minute interval. The G values obtained from these pictures were converted to their respective OD_600_ values using the manufacturer’s software, equation 1, and pre-calculated values of the constants (a=0.0234, b=1, c=1.09E-6, d=3.41, e=1.56E-13 and f=6.58).

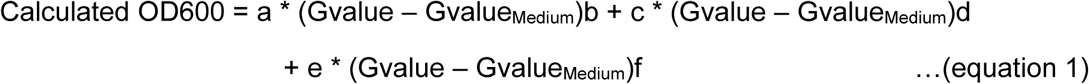

In some cases the estimates were clearly far too high due to outliers; Growth rates were truncated at 3 doubling times.h^-1^.

## Results

### Variation of the uptake of a lipophilic carbocyanine cation

3,3′-Dipropylthiadicarbocyanine iodide (IUPAC name (3-Propyl-2-{(1E,3E,5E)-5-(3-propyl-1,3-benzothiazol-2(3H)-ylidene)-1,3-pentadien-1-yl}-1,3-benzothiazol-3-ium iodide, Fig 1), commonly known as diS-C3(5) [108; 109], is a cationic carbocyanine dye that is accumulated by bacteria (both Gram-positive and Gram-negative) with energised membranes. Although culturability is the conventional metric for bacterial ‘viability’ [110; 111], diS-C3(5) has been exploited widely in microbiology to detect nominally ‘viable’ bacteria (at least those with intact membranes) in clinical, laboratory and environmental samples, especially using flow cytometry (e.g. [112-116]). Metabolising yeast cells can also accumulate it [117-121]. We note that the extent of fluorescence is not necessarily fully linear with concentration because of dye stacking [108; 109; 122](and see [123]) but is at least assumed to be more or less monotonic.

**Figure 1.**
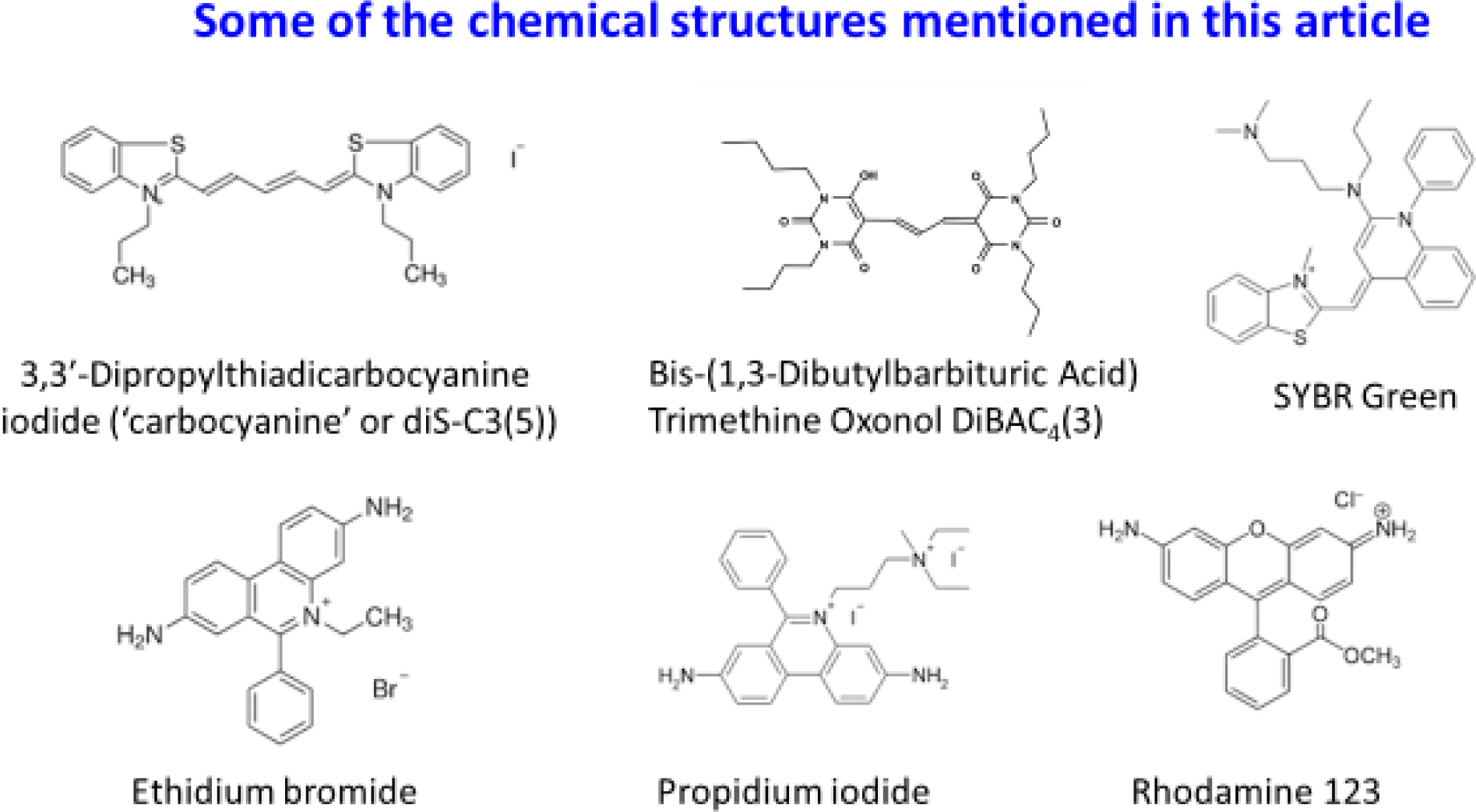
Chemical structures of some molecules described in this article.

The strategy used to assess diS-C3(5) uptake, gating on light scattering in ‘forward’ and ‘side’ directions by the bacteria and assessing uptake via red fluorescence (which is not interfered with by any autofluorescence) is precisely as was described previously [124]. It is obvious (Fig 2) that even in the wild-type strain there is considerable heterogeneity in the distribution of uptake between individual cells; this becomes even greater for the cells (ΔatpB) lacking the β subunit of the membrane ATP synthase. Fig 3 shows the mode fluorescence for the wild type strain and for knockouts of known transporter and certain other genes; the wild type and some of these are labelled in Fig 3. (Note that we have strong suspicions that the supposed tolC knockout was not in fact a knockout of tolC.) The very great breadth and complex shapes of the distributions is considered to be underpinned by the significant <u>number</u> of transporters involved, especially those driven by ATP, and their heterogeneous expression level distributions between individual cells [110].

**Figure 2.**
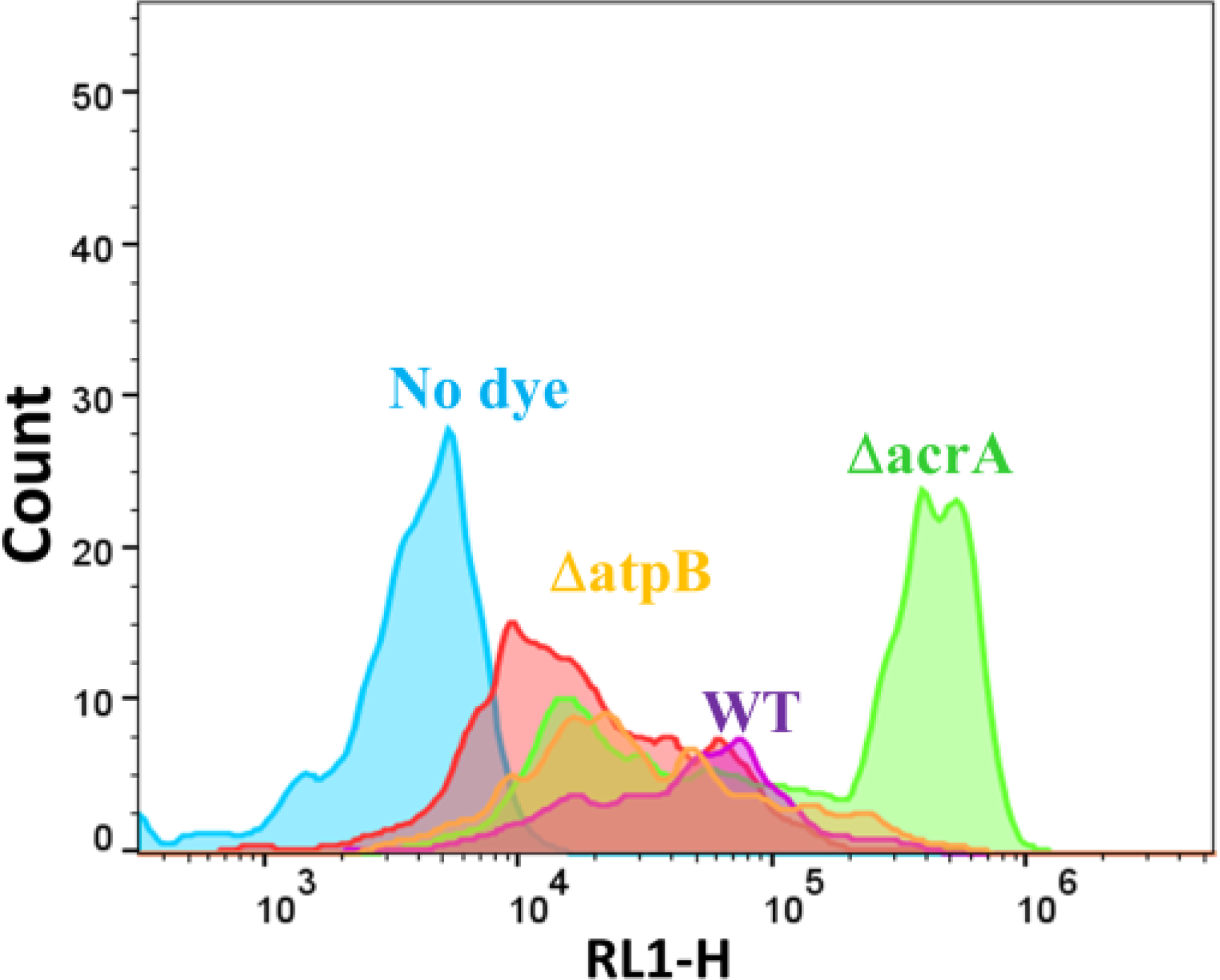
Typical cytograms of the wild-type strain (WT) stained (save for the no-dye control) with diS-C3(5) as described in Methods, along with other knockout strains. Those deleted in atpB or acrA are labelled with the relevant colours. Experiments were performed as described in the Materials and Methods section.

**Figure 3.**
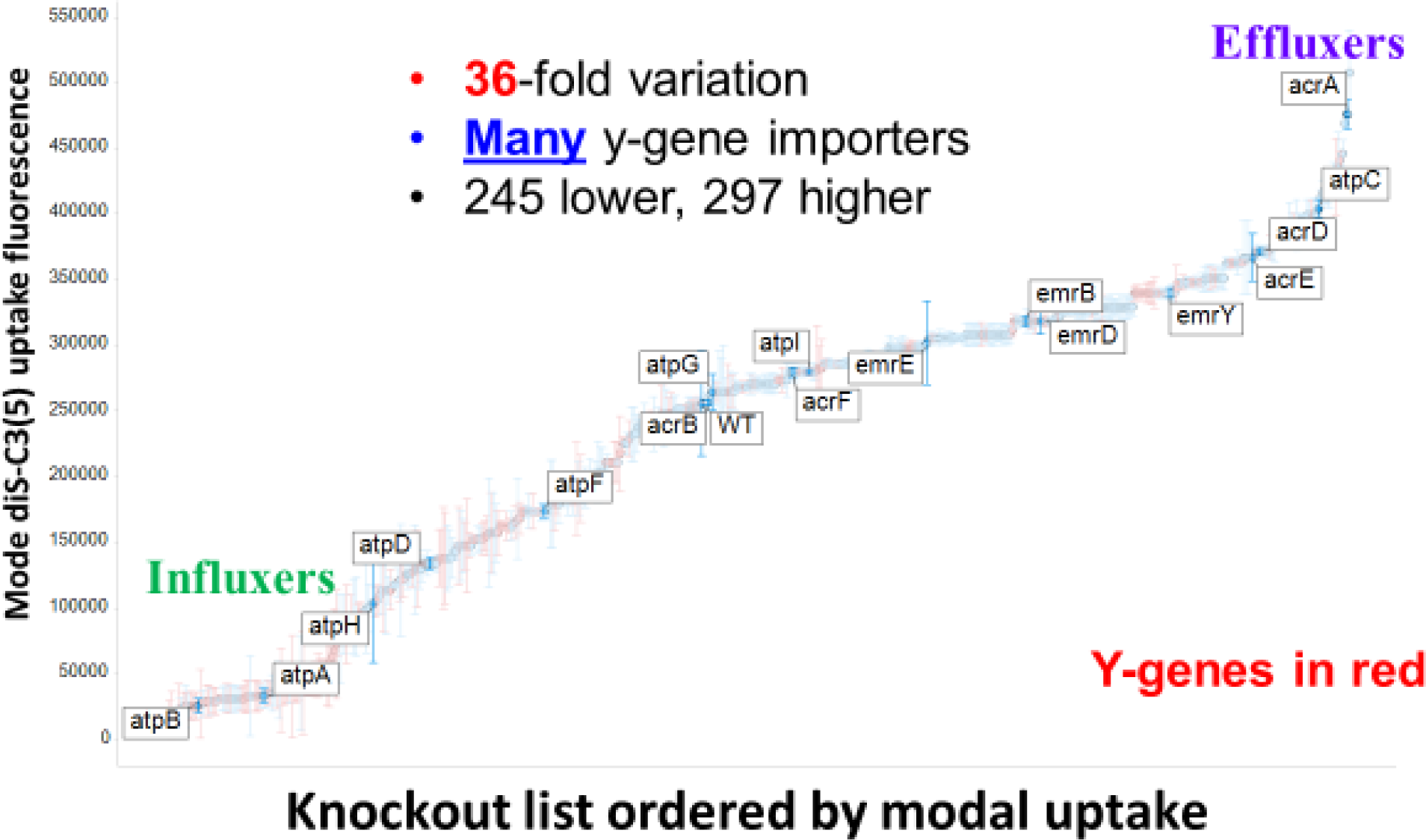
Variation in the mode uptake of diS-C3(5) into different knockout strains of the Keio collection as judged by flow cytometry. Experiments were performed as described in the legend to Fig 3. Experiments were performed as described in the Materials and Methods section.

The shape of the curve of Fig 3 is of interest, with two major break points, one after the lowest 75 and one before the highest 15 in terms of their effects on the steady-state uptake of DiS-C3(5). Several other important features emerge from this plot (Fig 3). (i) the range of steady-state uptake over the family of single-gene knockouts is very substantial, i.e. some 36-fold between the lowest and highest mode uptake, (ii) this range is somewhat asymmetric, in that the highest values are just two-fold greater than is that of the wild type (although we cannot exclude truncation due to fluorescence quenching [108]), while the lowest value is ∼18 times lower; (iii) the number of (y-)genes of unknown function that are uptake transporters (or whose removal most decreases the uptake of diS-C3(5)), encoded in red, is far greater than is the number of unknown effluxers (or those whose removal most increases its uptake; the three highest such y-genes are ygfQ, ydcS and ybbW, Table 1), (iv) the five highest values include four known drug effluxers (mdtJ, acrA, mdtI, mdtA), giving confidence in the strategy; (v) 297 knockouts are higher and 244 lower. If a very forgiving criterion of 50% above or below the WT is taken (fluorescence levels <128,000 or >384,000), we still have 115 knockouts below and 33 knockouts above these thresholds. While we do not know which ones are operating under each condition, this does imply the <u>potential</u> contribution of a very considerable number of transporters to the steady-state uptake of the dye, which is, of course, consistent with the very broad range of uptakes of an individual dye in a given cytogram (Fig 3).

**Table 1.** A selected subset of the most effective knockouts in terms of their ability to affect the uptake of diS-C3(5). Those whose knockout increased uptake are given in bold face.

| Gene | Comments | Uniprot ID | Representative reference(s) |
| --- | --- | --- | --- |
| *atpB* | F<sub>1</sub>-ATPase subunit | P0ABB0 | [130] |
| *atpA* | F<sub>1</sub>-ATPase subunit | P0AB98 | [130] |
| *ycdG (rutG)* | Predicted transporter/<br>putative pyrimidine<br>permease | W8ZH25/ P75892 | [131; 132] |
| *rbsB* | Ribose import binding<br>protein | P02925 | [133] |
| *ybiO* | ‘mechanosensitive ion<br>channel’ | P75783 | [134] |
| *yifK* | “probable transport<br>protein” (possibly amino<br>acid) | P27837 | [135] |
| *yliA/gsiA* | ATP-driven Glutathione<br>importer | P75796 | [136] |
| *ybiR* | Inner membrane protein | P75788 | (none) |
| *yccS* | Inner membrane protein | P75870 | (none) |
| *phoR* | Phosphate sensor<br>regulon | P08400 | [137] |
| *yejA* | Microcin<br>(?oligopeptide?)<br>importer | P33913 | [138] |
| ***mdtI*** | Spermidine export<br>protein | P69210 | [139; 140] |
| ***mdtJ*** | Spermidine export protein | P69212 | [139] |
| ***mdtL*** | Multidrug resistance protein (e.g. vs chloramphenicol) | P31462 | [139] |
| ***mdtA*** | Multidrug resistance protein (e.g. vs novobiocin) | P76397 | [141] |
| ***acrA*** | Multidrug efflux pump subunit | P0AE06 | [142] |
| ***ybbW*** | Putative allantoin permease | P75712 | [143; 144] |
| ***ydcS*** | Part of ABC transporter complex, possibly involved in poly-β-hydroxybutyrate synthesis/ polyamine efflux | P76108/A0A66RBW4 | [145] |
| ***ygfQ*** | Hypoxanthine permease | Q46817 | [146] |

Although it was not financially reasonable to determine any compensating or pleiotropic changes [125] at the level of the genome-wide transcriptome for these many hundreds of experiments, the simplest explanation for such data is that an increase in uptake following a knockout denotes the removal of an efflux transporter, while a decrease denotes that an influx transporter (or its source of free energy) has been knocked out. Either there are a massive number of pleiotropic effects on a smaller number of transporters or there are least 115 influx and 34 efflux transporters for diS-C3(5) (or both). Similarly, this vast (36-fold) range of uptake levels is not obviously consistent with the fact that uptake might reflect a membrane potential, although the activity of at least acrAB/tolC is in fact enhanced by an energised membrane. Indeed, the lowering of uptake in the ΔatpA/B knockouts, that would be expected to have a <u>higher</u> membrane potential [126], makes it absolutely clear that the uptake cannot mainly be driven by such a potential, and instead by ATP directly (generated by substrate-level phosphorylation). We note that a tolC knockout was not the highest (although it might be expected to inhibit many RND-type efflux pump activities), but it seems that this strain may have had a mutation elsewhere as tolC was still present as judged by colony PCR and our own attempts to remove it were unsuccessful. Similar suspicion applies to acrB (Fig 3, which would be anticipated to lie near acA). But the general principle remains clear: individual gen knockouts lead to a huge range the steady-state uptake of the dye that cannot be ascribed to changes in a membrane potential with which its distribution might equilibrate.

To confirm these general findings, we took a few of the gene designations whose removal in the Keio collection led to the greatest or lowest uptake, and assessed their behaviours in the corresponding strains in the ASKA collection of overexpressed genes. As anticipated, overexpression of these effluxers lowered the uptake levels to less than that of the wildtype, while overexpression of putative uptake transporters raised the steady-state level of their uptake considerably (Fig 4), albeit not to the levels of the highest effluxer KOs, implying some spare kinetic capacity in the latter.

**Figure 4.**
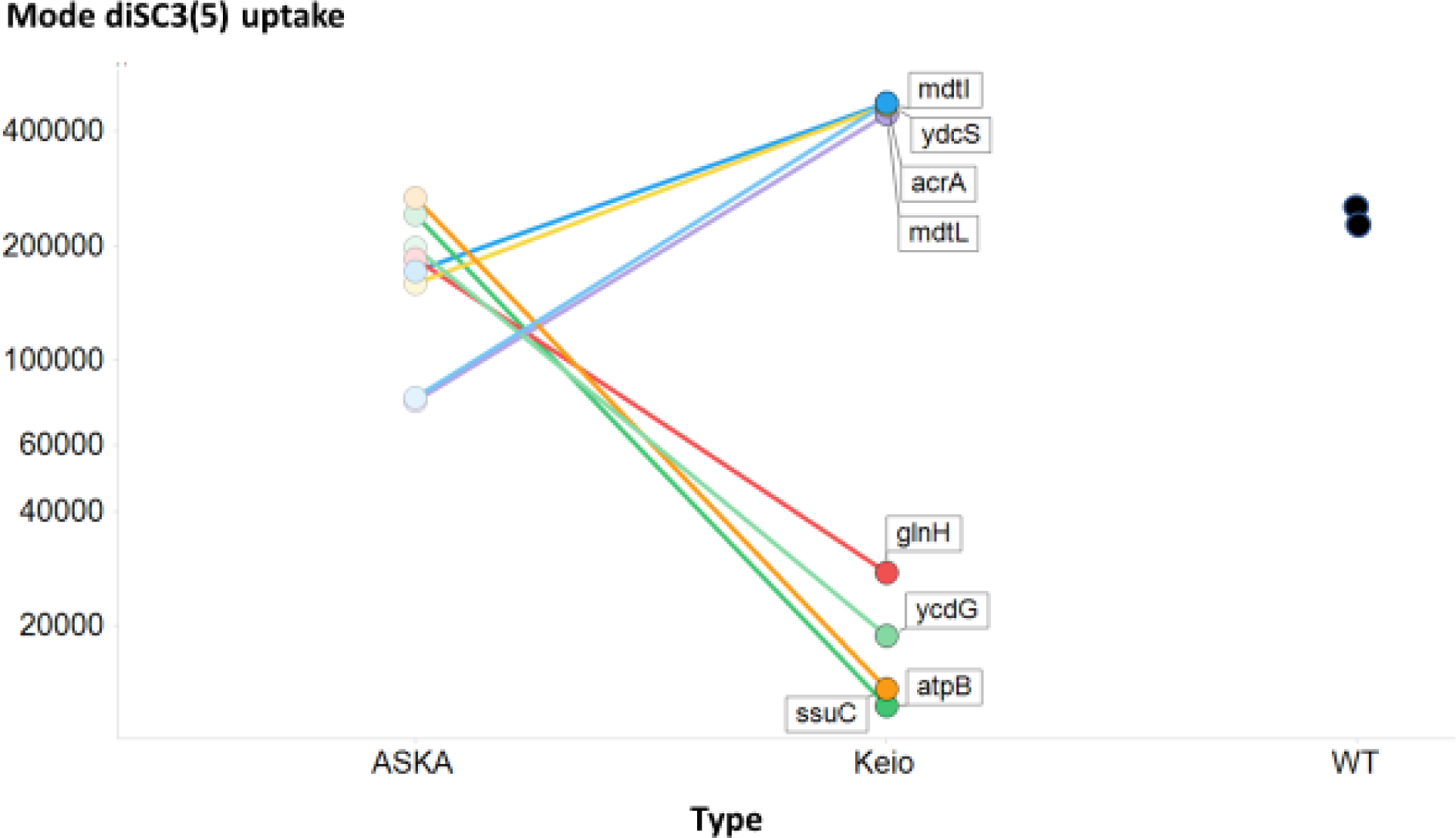
Effect of overexpression and knockout of a series of genes whose knockout causes major changes (both up and down) in the modal uptake of diSC3(5) relative to that of the wild type. Experiments were performed as described in the Materials and Methods section.

To analyse more closely the main conclusion of this section – that the steady-state uptake level of the diS-C3(5) molecule is determined significantly by multiple proteins including transporters – we highlight some of the most significant knockouts in Table 1. A number are unsurprising, which gives confidence in the idea that our basic method is sound. Among the most potent of the non-transporters at inhibiting uptake were knockouts of the genes encoding the two main (α and β) subunits of the membrane ATP synthase, implying strongly that the uptake transporters were powered directly by ATP, rather than say by an electron-transport-derived energised membrane (often referred to as a protonmotive force). This is consistent with the relative prevalence of ATP-driven transporters in prokaryotes [35].

The complement of y-genes of unknown or poorly annotated function stands at around 35% of the total, with transporters being over-represented among them [127]. It is of particular interest that so many y-genes are represented among the KOs showing the biggest effects; clearly we have much to learn, as with mammalian cell transporters of pharmaceutical drugs [128; 129], about their ‘natural’ substrates.

It is of course entirely arbitrary to pick the top few only (all are given in the Supplementary Information), and we would add that one set of interest is represented by potC and potE (as well as other products of *pot* genes) that are components of a (cationic) sperm(id)ine import/efflux system [147-150]. Obviously it is entirely reasonable that this might serve, when active, to remove a cation such as diS-C3(5), and the fact that the potent effluxers mdtI and mdtJ are also spermi(di)ne effluxers (Table 1) lends weight to this view. In a similar vein, the KOs of mdtH and mdtK [140] have a significantly reduced uptake; although they were originally tagged as effluxers (in the sense of multidrug transporters), it seems more likely that they are in fact antiporters, normally contributing to the uptake (here) of the cyanine dye.

### SYBR Green

SYBR Green is another cationic dye, and increases its fluorescence massively on binding to (especially) double-stranded DNA [10; 151]. It is considered to be ‘membrane-permeable’ (by whatever means) and it is widely used both in environmental and general microbiology (e.g. [10; 151-155]) and in mammalian cell biology (e.g. [156]). SYBR Green also contains a cyanine motif (Fig 1), and using our standard methods of cheminformatics [129; 157; 158], we noted that SYBR Green and DiS-C3(5) have a Tanimoto similarity to each other of 0.731 when encoded using the RDKit (www.rdkit.org) “patterned” fingerprint.

A similar experiment to that performed with diS-C3(5) was performed with SYBR Green. Typical cytograms of SYBR Green uptake are shown in Fig 5, and the full set of results (slightly fewer KOs than for diSC3(5)) shown in Fig 6. Again there is a huge range of uptakes (69-fold), the lowest being some five-fold lower and the largest some thirteen-fold greater than that of the wild type. Because of the greater range, and the occasionally bimodal peaks such as that for ΔatpG in Fig 5, the median fluorescence values are given on a logarithmic scale. All data are again given in Supplementary Table 1. Some of this variation can of course reflect differences in the DNA content of the cells, since this is a function of the earlier growth rate [159; 160], though this typically does not exceed about 8 chromosome contents even at the fastest growth rates [107; 161], so variations in DNA content alone could not conceivably explain this range. We therefore also performed growth rate experiments for almost all the strains tested; there was no correlation between the uptake of SYBR green and either the growth rate (Fig 7, r^2^ ∼0.02) or the stationary phase OD (Fig 8, r^2^ ∼0.015), indicating clearly that variations in DNA content were not a significant contributor to these findings in intact cells.

**Figure 5.**
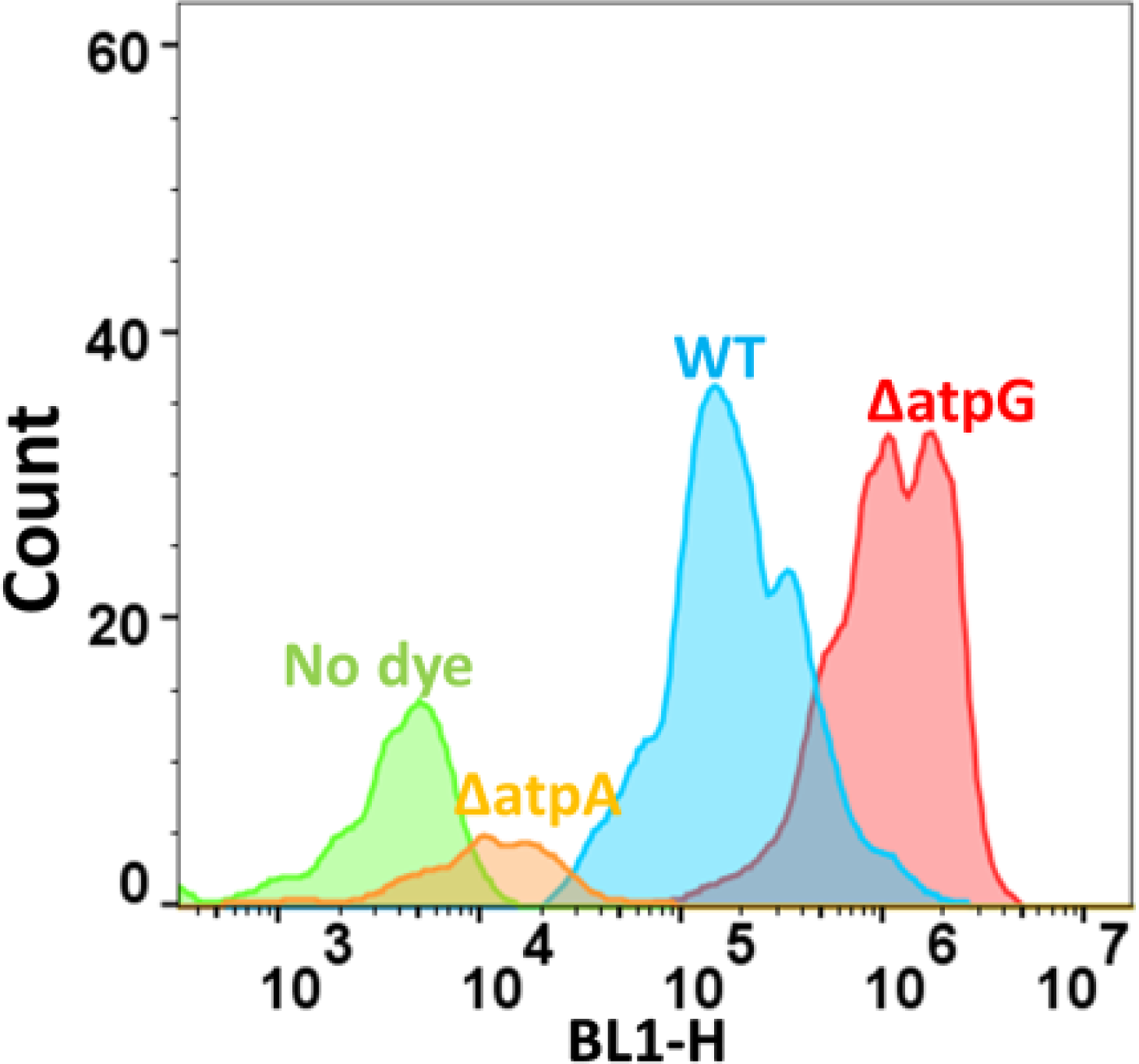
Typical cytograms of the uptake of SYBR Green in the wild-type strain (WT) and some other, knockout strains stained with SYBR Green as described in Methods (save for the no-dye control), along with other knockout strains.

**Figure 6.**
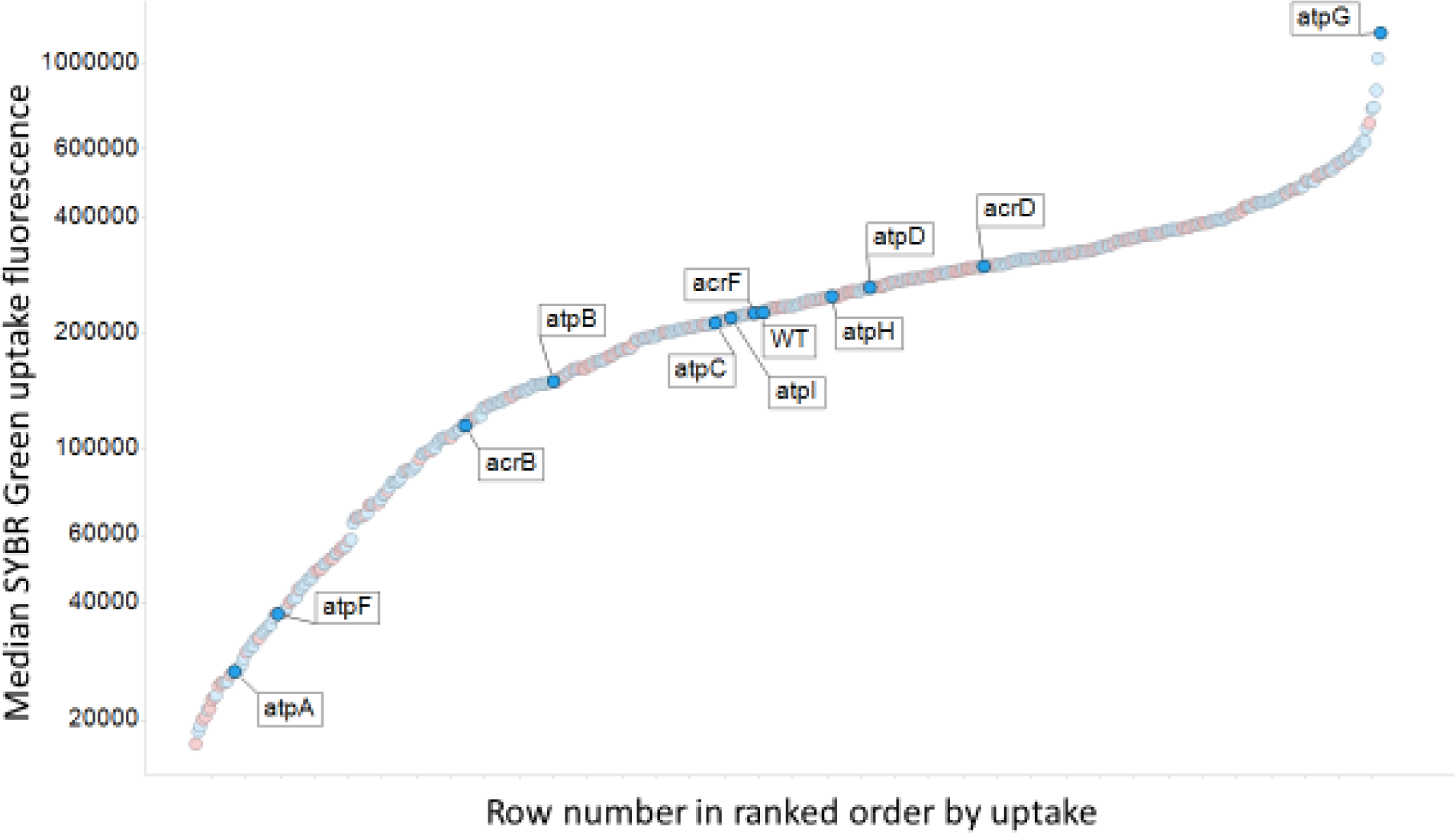
Median uptake in ranked order of SYBR Green uptake into E. coli single-gne knockout strains, with a small subset of gene names marked. Y-genes are encoded in red. Experiments were performed as described in the Materials and Methods section

**Figure 7.**
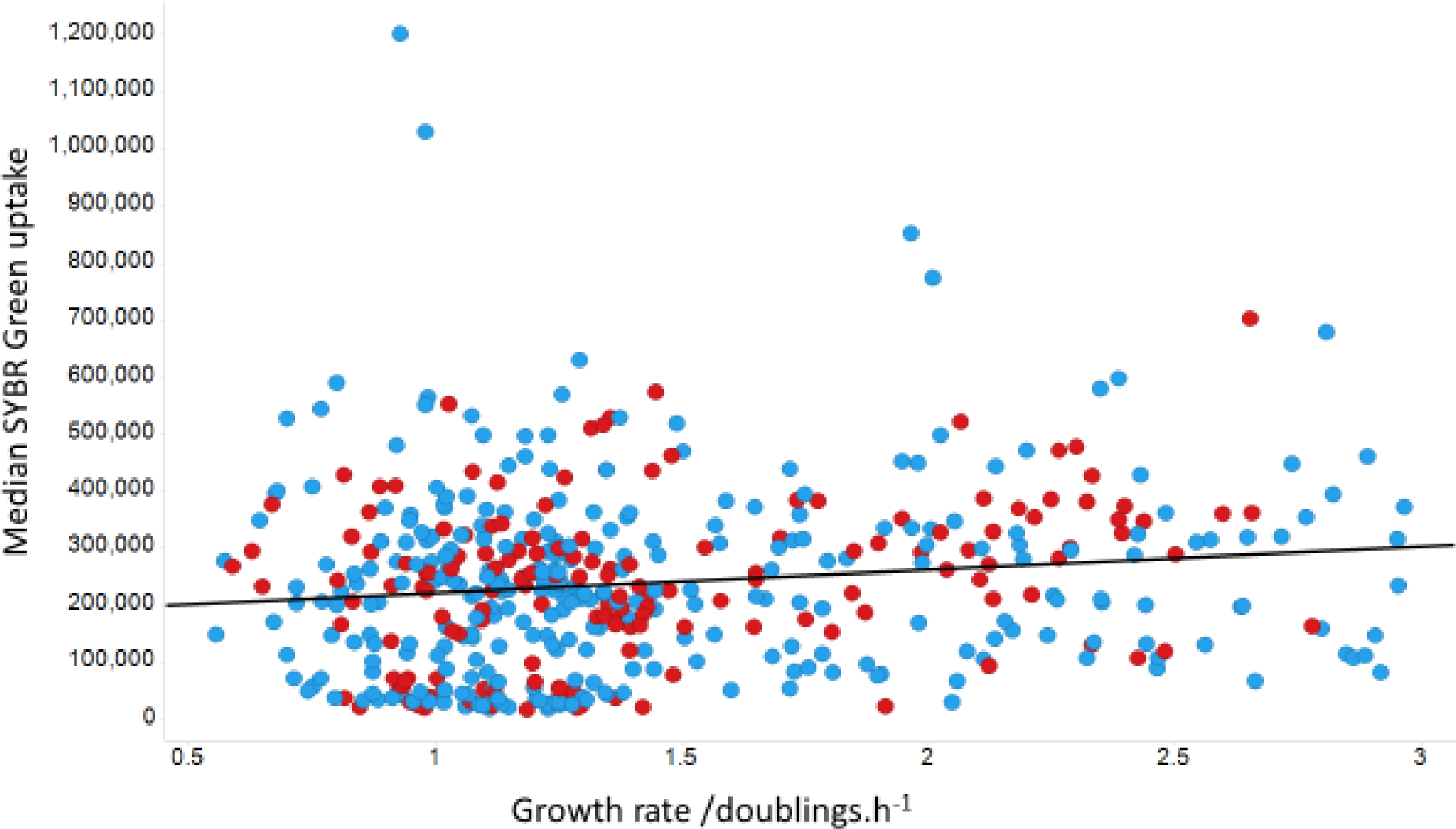
Lack of relationship between the median extent of uptake of SYBR Green and growth rate. Y-genes are encoded in red. Experiments were performed as described in the Materials and Methods section.

**Figure 8.**
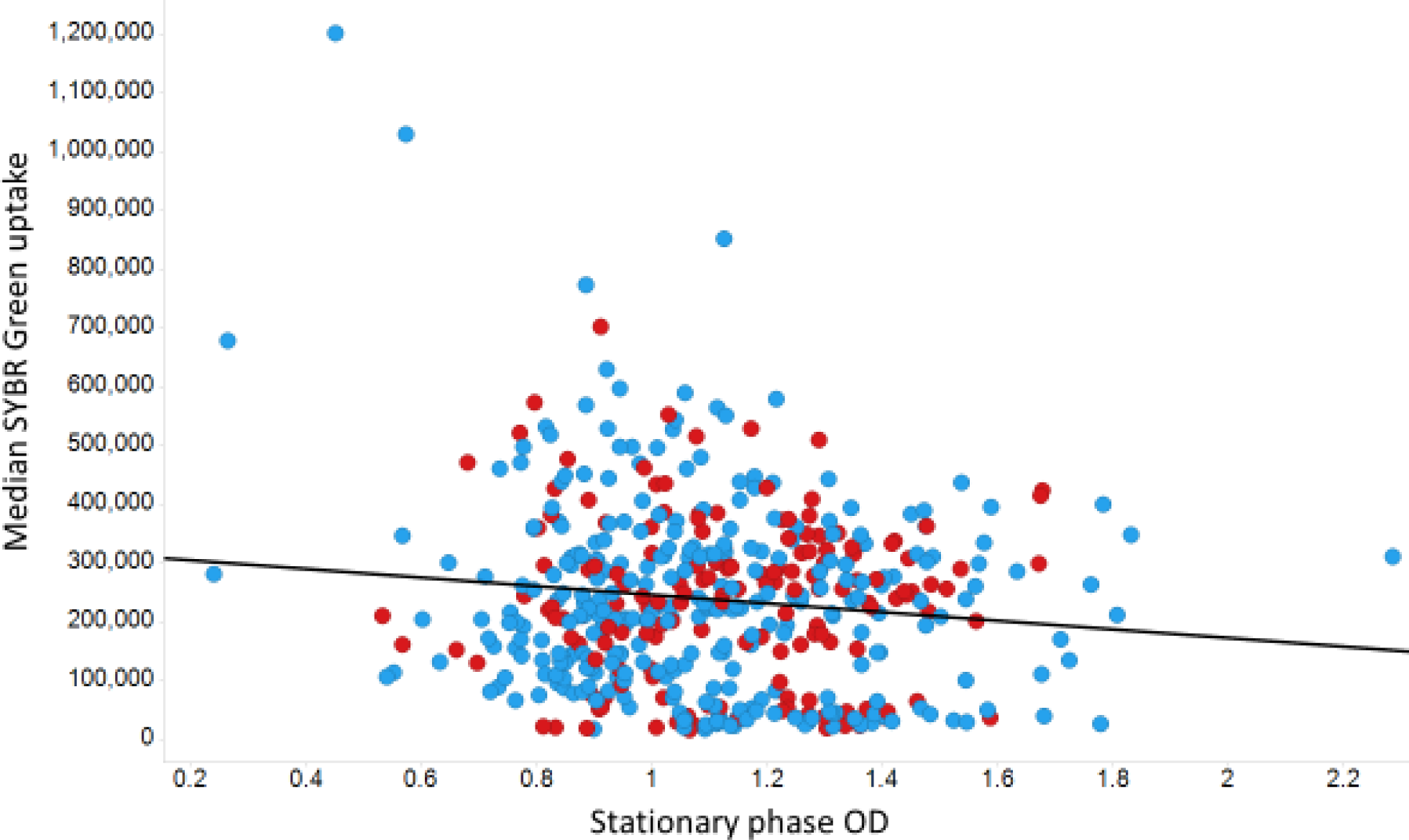
Lack of relationship between the median extent of uptake of SYBR Green and stationary phase OD. As before, y-genes are encoded in red. Experiments were performed as described in the Materials and Methods section.

By contrast with the diS-C3(5) data (Fig 3), there are considerable differences (Fig 9), in that the phoR knockout now shows the second greatest uptake (rather than the eighth lowest). Again there is a considerable preponderance of y-genes in the part of the figure where their removal lowers uptake relative to the wild type. A subset of genes with the ‘highest’ and ‘lowest’ effects is given in Table 2.

**Table 2.**
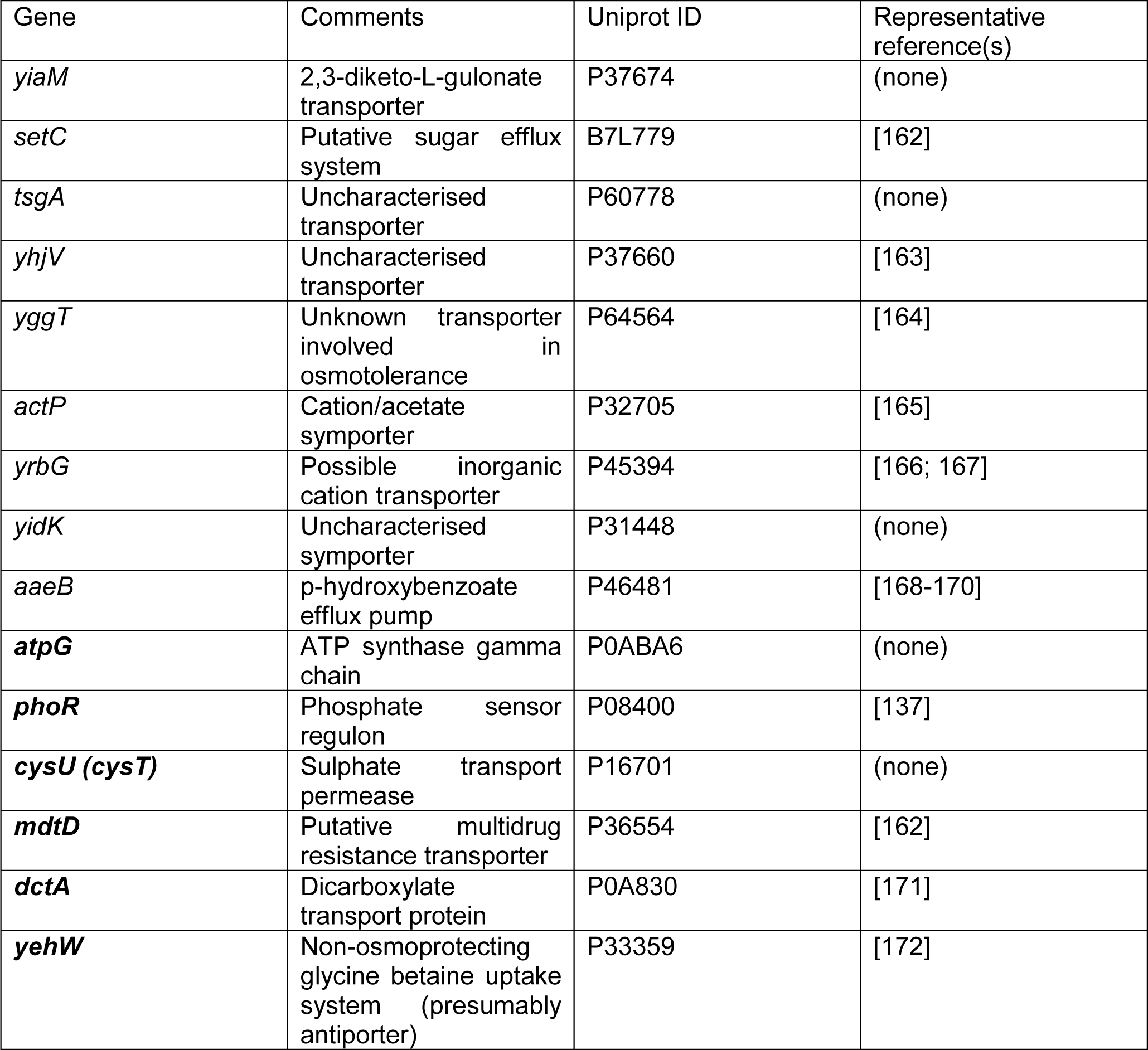
A selected subset of the most effective knockouts in terms of their ability to affect the uptake of diS-C3(5). Those whose knockout increased uptake are given in bold face.

**Figure 9.**
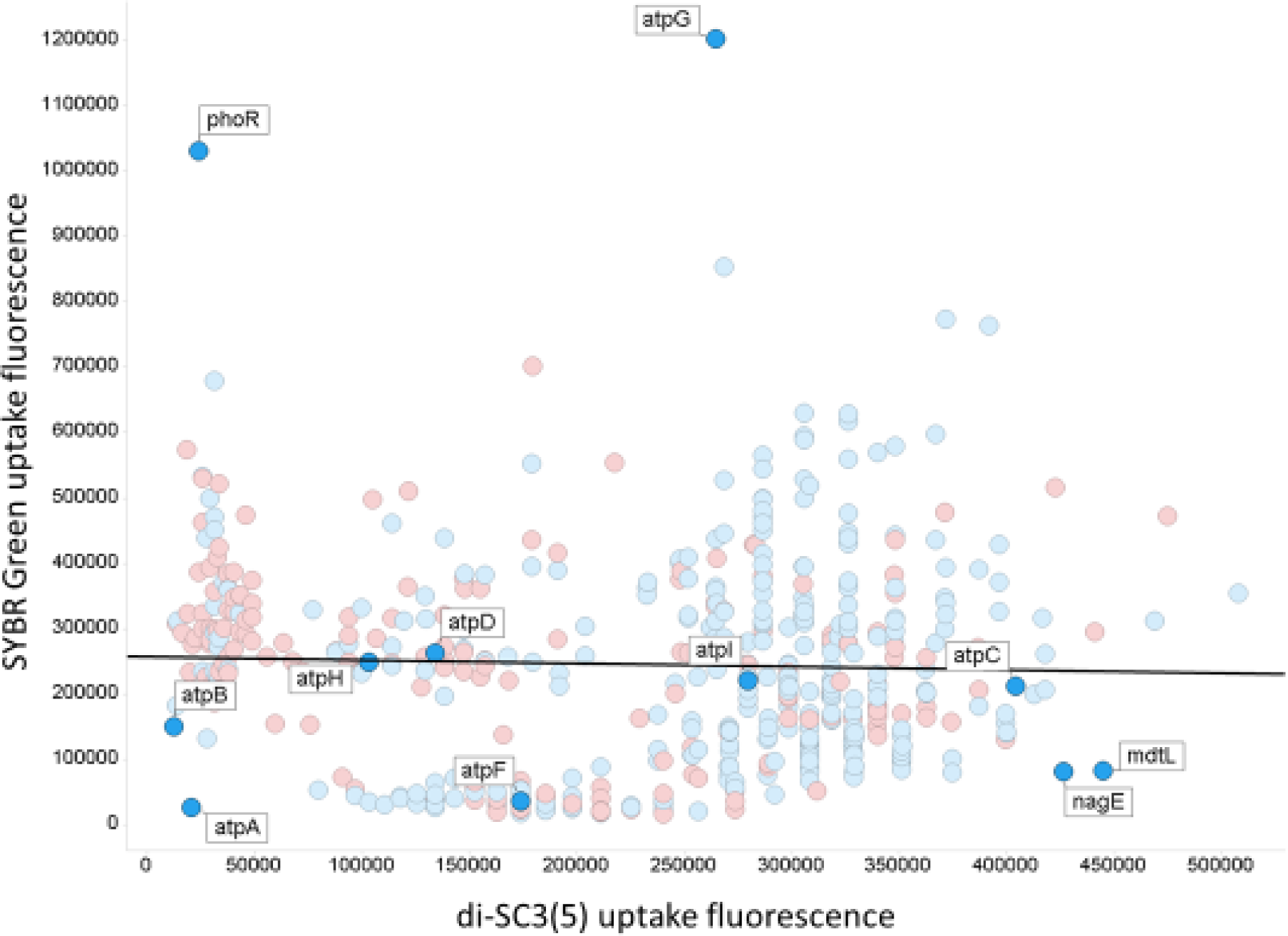
Relationship between The uptake of SYBR Green (ordinate) and of diS-C3(5) (abscissa. The line is a line of best fit (r^2^∼0.002). Also shown are some of the knockout labels as discussed in the text. Symbols for Y-genes are in red, others in blue. Experiments were performed as described in the Materials and Methods section.

Despite the fact that both dyes contain a cyanine (benzimidazole) motif, there is absolutely no correlation (r^2^ ∼ 0.002) between the uptake of the two dyes in individual knockouts (Fig 6). Thus, mdtL and nagE are among the highest for diS-C3(5) but among the lowest for SYBR Green, while the converse is true for phoR, zntA (zinc/lead-transportng ATPase) and yifK (‘probable amino acid transporter protein’). Such findings imply that some transporters have a fairly unusual specificity, even if they are labelled (as is mdtL) as multidrug resistance proteins. In the case of SYBR Green, the heterogeneity in uptake can clearly contribute to the heterogeneity in staining observed [10] when cells are not permeabilised.

It was also of interest to assess the effect of chemical inhibitors on the ability of transporter knockouts to take up SYBR Green. In contrast to earlier work [173] where we studied effects on gene transcription, the interest here was in direct, acute effects. Table 3 shows the effects of a few such molecules, that of chlorpromazine, a known inhibitor of acrAB activity [174-176], and clozapine (a second-generation anti-psychotic) [177] being particularly striking. Note (see Methods) that different conditions were used from those of Figs 2-6. The very large increase upon chlorpromazine addition (more than 25-fold) might be taken to imply that it can inhibit multiple effluxers, and suggests that it might be a useful adjunct therapy in cases of antimicrobial resistance.

**Table 3.** Effects of various modifiers on steady-state uptake of diSC3(5) and SYBR Green by wild-type *E. coli.* WT means no inhibitors added

| S.No. | Efflux Inhibitors | diS-C3(5) | SYBR Green I |
| --- | --- | --- | --- |
| 1 | Rifampicin | 94 | 62.3 |
| 2 | Propenacid | 60.8 | 58.4 |
| 3 | Elacridar | 61 | 63.8 |
| 4 | Verapamil | 56.4 | 59.5 |
| 5 | MK571 | 33.1 | 40.3 |
| 6 | KO143 | 51.7 | 59.1 |
| 7 | Mefloquine | 37.6 | 93 |
| 8 | Cyclosporin A | 64.3 | 58.1 |
| 9 | Chlorpromazine | 54.9 | 1347.5 |
| 10 | Olanzapine | 128.9 | 106.6 |
| 11 | Lapatinib | 43.2 | 59.5 |
| 12 | Tariquidar | 50.4 | 59.1 |
| 13 | Clozapine | 33.4 | 350.8 |
|  | WT | 51 | 66.2 |

## Discussion

There is present lively debate as to whether a majority of xenobiotic uptake through cellular membranes occurs via whatever phospholipid bilayer may be present, or whether Phospholipid Bilayer diffusion Is Negligible (a view referred to as “PBIN”; [27; 178]). In this latter view, it is recognised that potentially a great many transporters can and do interact with a given molecule [27; 178] (see also [29]). This would be unsurprising given that the typical <u>known</u> numbers of <u>binding targets</u> for pharmaceutical drugs is ∼six [26; 179].The present data are entirely consistent with (indeed lend strong support to) this view.

If a closed biological cell or organelle maintains a transmembrane electrical potential difference ΔΨ relative to that of its adjacent, external phase, it is in principle possible to estimate that potential by allowing a freely membrane-permeable (usually lipophilic) ion to come into a Nernstian equilibrium with it (e.g. [180-189]). For a negative-inside potential, and concentrative uptake by a lipophilic cation such as methyltriphenylphosphonium (TPMP^+^), ΔΨ may be related to the ratio of internal a_in_ and external a_out_ activities of the ions by:

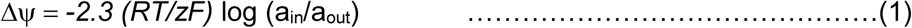

where T is the absolute temperature, R the universal gas constant, F Faraday’s constant, and z the charge on the lipophilic cation. RT/F is about 60 mV at room temperature, such that a ΔΨ of 60 mV equates to a concentration ratio for a monovalent cation of 10:1, a ΔΨ of 120 mV a concentration ratio of 100, and so on. It is often assumed that the uptake of such lipophilic cationic dyes by bacteria reflects the existence and magnitude of such a transmembrane potential. However, import and export transporters invalidate such estimations of ΔΨ, their activities respectively causing an over- and under-estimate, and this has been shown clearly to occur for the uptake of dibenzyldimethylammonium in baker’s yeast [70] and of Tl^+^ in bacteria [190-192]. Confidence in any such estimations of ΔΨ is bolstered if the equilibrium uptake ratio is independent not only of the concentration but the nature of the lipophilic cations employed, and also of any anions that may be present. In practice, and although such difficulties are commonly ignored, these requirements are rarely if ever met [180; 182; 193; 194].

However, in the present work we have shown clearly that a variety of cations enter cells via a large number of transporters, especially those driven by ATP directly, so whatever their uptake is reflecting it cannot simply be a bulk transmembrane potential. Indeed, the loss of ATP synthesis by electron transport-linked phosphorylation in the ΔatpA and ΔatpB strains might be expected to <u>increase</u> the level of any such membrane potential [126] but instead the extent of uptake is substantially reduced. This is very strong evidence against any such equilibration of uptake with a membrane potential. In the case of SYBR Green, it also calls into question the use of that dye for estimating quantitatively the amount of DNA in live, non-permeabilised cells, and (using growth rate as a surrogate [159; 195] for DNA content) no such relationship was found.

By contrast, a possible benefit of our findings, for those interested in estimating transporter activities, is that if overexpression of a particular transporter causes most of the uptake (or efflux) flux to occur via it (as in Fig 4), competition or trans-stimulation assays with a fluorophore provide a powerful and potentially high-throughput [44; 196] method for measuring QSARs. The fact that the substrate specificities of individual transporters are typically rather different from each other (Fig 6) implies that there could indeed be much value in pursuing this more widely.

## Acknowledgments

We thank the BBSRC (grants BB/R000093/1 and BB/P009042/1) and the Novo Nordisk Foundation (grant NNF10CC1016517) for financial support.

